## Supplemental Material for "Universal microbial reworking of dissolved organic matter along environmental gradients"

This .pdf file includes:
Supplementary Figures 1 - 3
Supplementary Tables 1 - 10
Supplementary Methods
Supplementary References

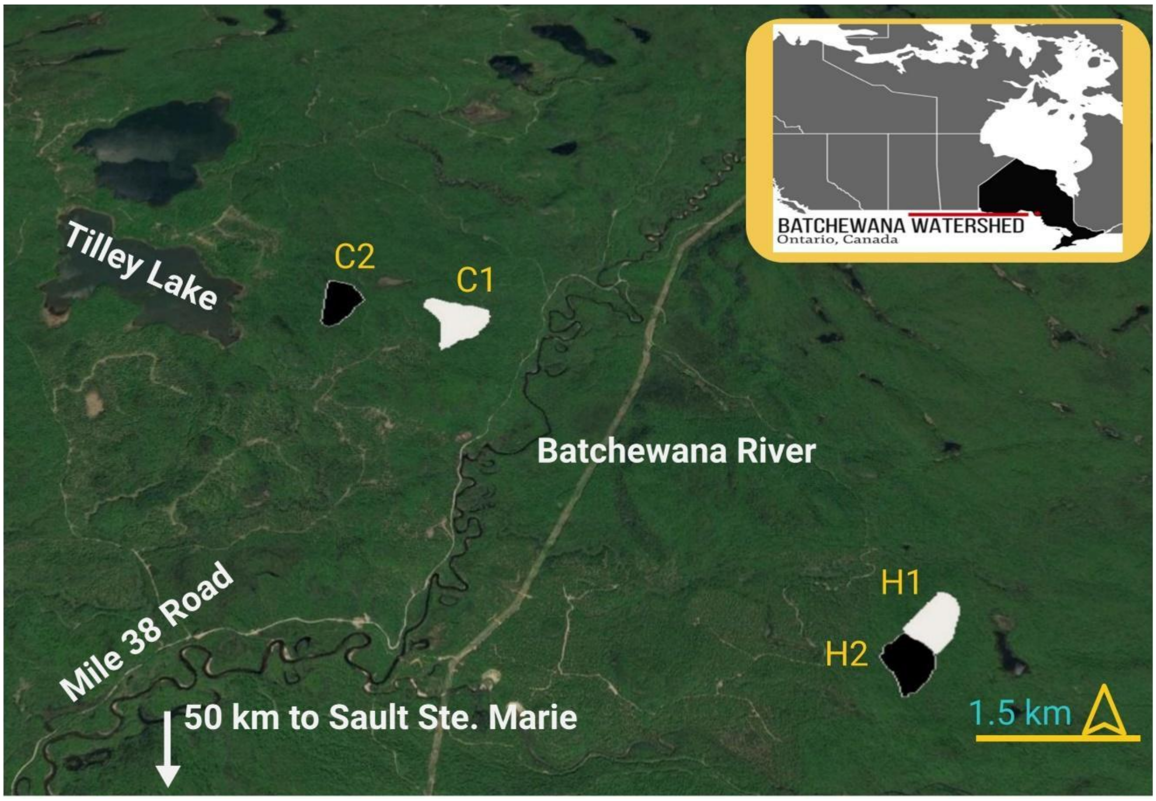


**Figure S1: Study area within Batchewana watershed, central Ontario, Canada with four replicate catchments named C1, C2, H1, H2.** These catchments were recruited as part of a forest harvest experiment and the names reflect this design: C = control and H = harvest. All samples were taken prior to these catchments being harvested and therefore can be treated as replicates. All geographic coordinates for sampling points can be accessed and downloaded from: https://www.gaiagps.com/public/VKMlEPlvf1t6ownd6Vulfm8T.

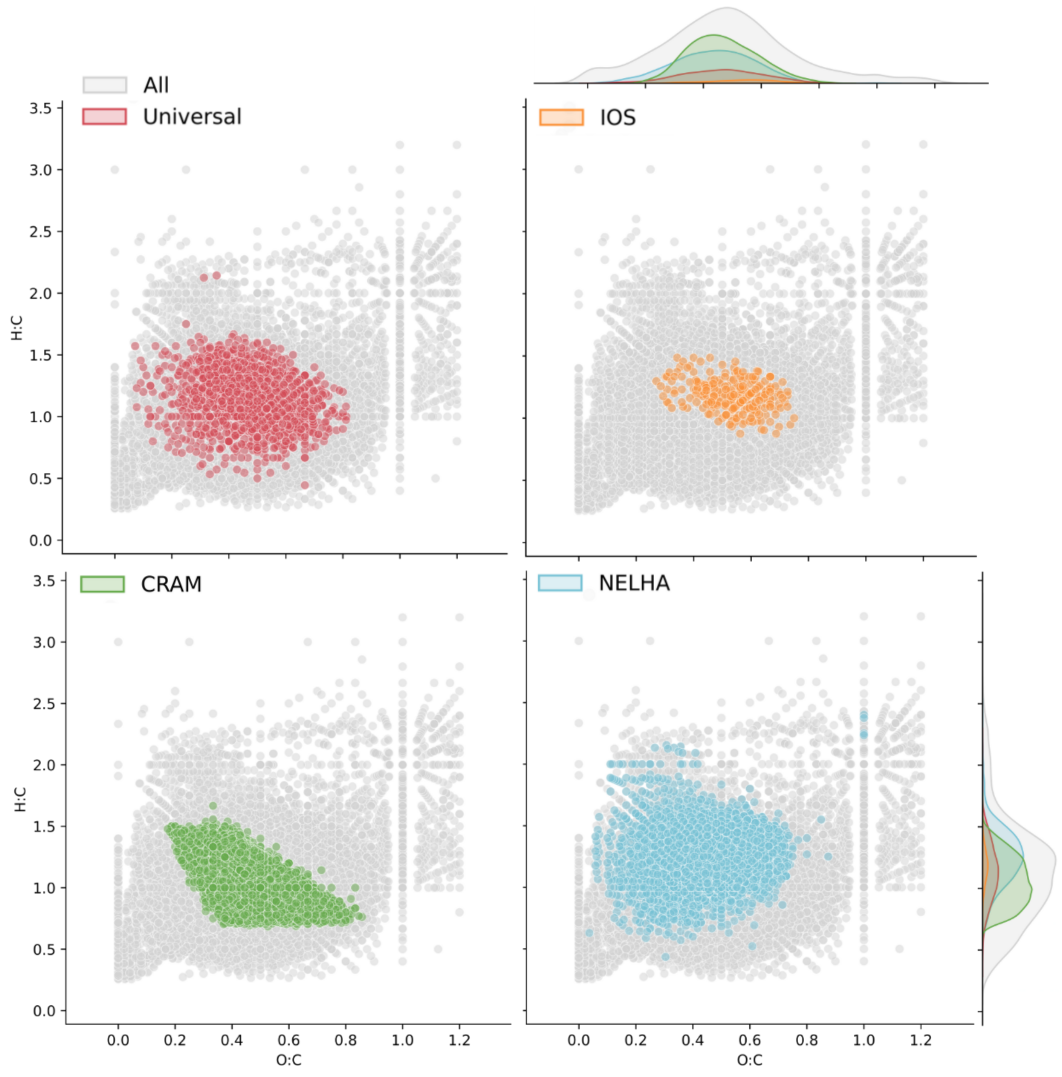


**Figure S2: Molecular properties are shared by different definitions of degradation end-products that should be universally distributed.** Each point is the elemental ratio (hydrogen to carbon and oxygen to carbon) of a molecular formula detected in our study ("All" category, n=9327). We further coloured these points according to whether they were present in every soil and stream sample in this study ("Universal"; n=1216), carboxyl-rich alicyclic molecules (CRAM, n = 3735), found in a subset of molecular formulae representing aged, degradation end-products identified in marine environments[1,2] (IOS, n=233), or found in a deep-sea reference sample from the Natural Energy Laboratory of Hawaii Authority (NEHLA) facility and expected to be dominated by degradation end-products (n=3019). CRAM were defined as molecular formulae that contained double bond equivalent (DBE) to C ratios of 0.30 to 0.68, DBE to H ratios of 0.20 to 0.95, and DBE to O ratios of 0.77 to 1.75[3]. Marginal axes show density curves for each compound category.

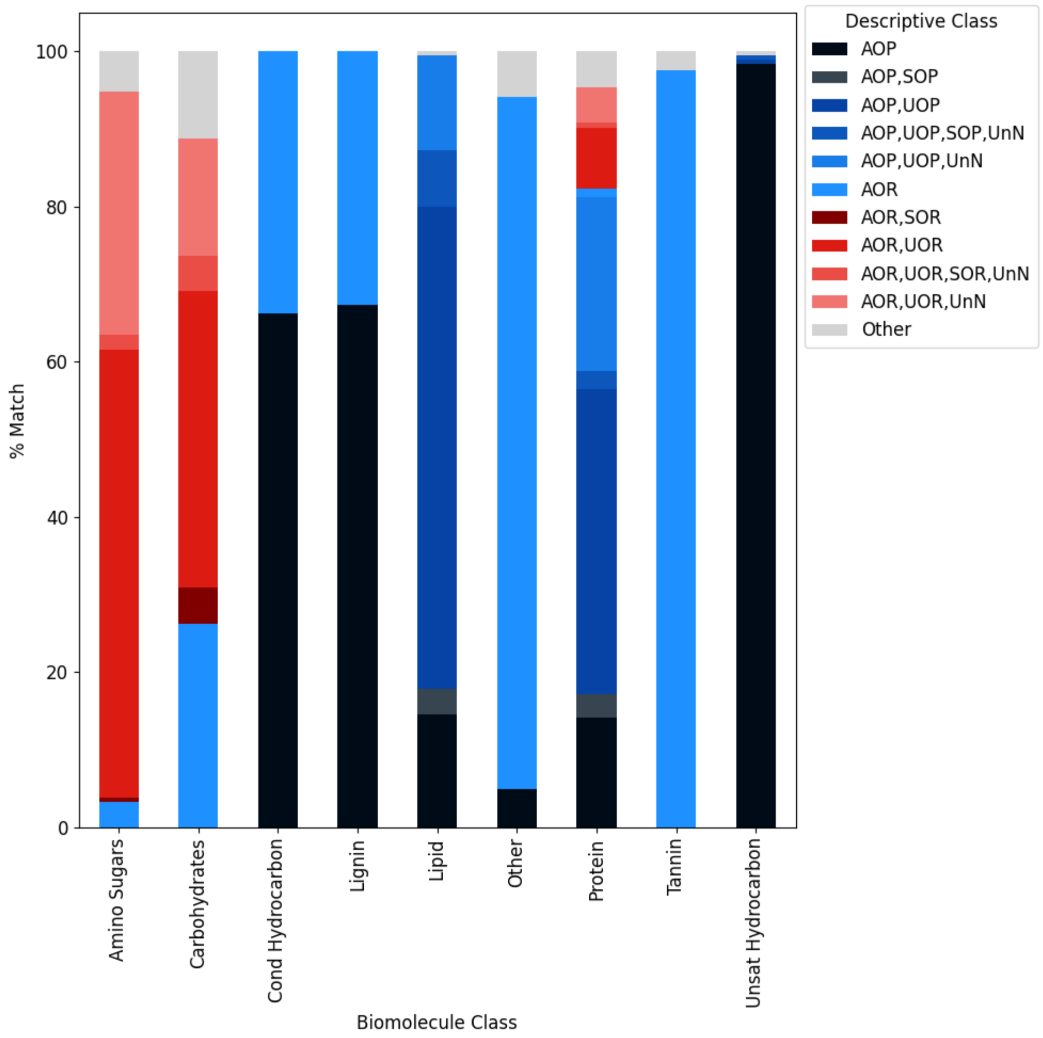


**Figure S3: Comparison of compound classifications.** We reclassified each molecular formula assigned to a compound class in the Main Text after ref. [4] using the descriptive classes of ref. [5]. Ref. [5] incorporates additional information on the modified aromaticity index and double bond equivalents of each formula in addition to the H:C and O:C ratios used by ref. [4]. The resulting descriptive classes can overlap and include: AOP = aromatic, oxygen-poor, SOP = saturated oxygen-poor, UOP = unsaturated oxygen-poor, UnN= unsaturated with N, SOR= saturated oxygen-poor or Other = unclassified.

**Table S1: Linear models predicting the number and relative abundance of universal compounds.** We modelled the proportion of molecular formulae that were counted in every sample (n=1216), along with their relative abundance (i.e. summed normalised signal intensity). We compared these with the proportion of molecular formulae and relative abundance that overlapped with the "Island of Stability" (IOS, n=233) – a literature-based definition of universal-like molecular formulae[1,2,6] (Fig. S2). We also modelled the relative abundance of compounds in our samples that could be defined as carboxyl-rich alicyclic molecules (CRAM), which contain double bond equivalent (DBE) to C ratios of 0.30 to 0.68, DBE to H ratios of 0.20 to 0.95, and DBE to O ratios of 0.77 to 1.75[3] (n=3735), and compounds found in a deep-sea reference (n=3019) collected near the Natural Energy Laboratory of Hawaii Authority (NEHLA) facility, which should be dominated by widely distributed, degradation end-products. Values in cells are mean estimated effects ± standard error relative to the intercept, with the intercept expressed relative to zero. Proportions were modelled using a binomial error structure, so effects are expressed on a logit-scale and correspond with odds ratios. Bolded values were statistically significant at ***$p < 0.001$, **$p < 0.01$, *$p < 0.05$. Adjusted $R^2$ for linear models with binomial error structures were estimated using the MuMIn package in R[7]. For all, degrees of freedom = 67.

| *Coefficient* | **Proportion of All Compounds** | **Abundance of All Compounds** | **Proportion IOS** | **Abundance of IOS** | **Abundance of CRAM** | **Abundance of NELHA** |
|---|---|---|---|---|---|---|
| Intercept: Site C1, 5 cm depth, shoulder position | **0.26 *** (0.03)** | **0.59 ** (0.06)** | **0.04 *** (0.00)** | **0.20 *** (0.02)** | **0.58 *** (0.04)** | **0.62 ** (0.11)** |
| Site C2 | **1.19 * (0.10)** | 0.53 (0.04) | **1.14 * (0.07)** | 0.51 (0.03) | 0.51 (0.02) | 0.53 (0.06) |
| Site H1 | 1.12 (0.09) | 0.51 (0.03) | 1.09 (0.07) | 0.52 (0.03) | **0.52 * (0.02)** | 0.54 (0.06) |
| Site H2 | 1.05 (0.09) | 0.50 (0.03) | 1.05 (0.07) | 0.52 (0.03) | **0.52 * (0.02)** | 0.54 (0.06) |
| 15 cm | 1.00 (0.08) | 0.51 (0.04) | 1.00 (0.06) | 0.52 (0.03) | 0.52 (0.02) | 0.54 (0.06) |
| 30 cm | 1.14 (0.10) | **0.57 *** (0.04)** | 1.11 (0.07) | **0.59 *** (0.04)** | **0.56 *** (0.03)** | **0.65 *** (0.07)** |
| 60 cm | **1.19 * (0.10)** | **0.60 *** (0.04)** | **1.14 * (0.07)** | **0.63 *** (0.04)** | **0.59 *** (0.03)** | **0.67 *** (0.07)** |
| Backslope | 1.01 (0.09) | 0.51 (0.04) | 1.01 (0.07) | 0.52 (0.03) | 0.52 (0.02) | 0.53 (0.06) |
| Footslope | 0.99 (0.08) | 0.51 (0.04) | 0.99 (0.06) | **0.55 ** (0.04)** | **0.55 *** (0.03)** | **0.59 ** (0.06)** |
| Toeslope | 1.16 (0.10) | **0.55 ** (0.04)** | 1.12 (0.07) | **0.57 *** (0.04)** | **0.57 *** (0.03)** | **0.63 *** (0.07)** |
| Stream | 0.90 (0.13) | **0.43 * (0.05)** | 0.92 (0.10) | **0.38 *** (0.04)** | **0.40 *** (0.03)** | **0.33 *** (0.06)** |
| $R^2$ | 0.20 | 0.42 | 0.21 | 0.59 | 0.64 | 0.54 |

**Table S2: Linear models predicting the molecular mass of DOM.** Molecular mass (Da) is equivalent to the mass to charge ratio (m/z) because all molecules are singly charged. Values in cells are mean estimated effects ± standard error relative to the intercept, with the intercept expressed relative to zero. Bolded values were statistically significant at ***p < 0.001, **p < 0.01, *p < 0.05. For all, degrees of freedom = 67.

| *Coefficient* | **Intensity-Weighted Mass (All Compounds)** | **Mass (All Compounds)** | **Mass (Non-Universal Compounds)** |
|---|---|---|---|
| Intercept: Site C1, 5 cm depth, shoulder position | **437.94 *** (8.15)** | **476.77 *** (8.33)** | **491.24 *** (9.62)** |
| Site C2 | -4.27 (5.16) | -7.07 (5.27) | -7.02 (6.08) |
| Site H1 | -0.37 (5.16) | -7.19 (5.27) | -7.88 (6.08) |
| Site H2 | 1.47 (5.16) | -4.29 (5.27) | -4.32 (6.08) |
| 15 cm | 4.04 (5.31) | -1.03 (5.43) | -1.62 (6.27) |
| 30 cm | -1.60 (5.31) | **-11.63 * (5.43)** | **-13.68 * (6.27)** |
| 60 cm | 10.70 * (5.31) | -3.53 (5.43) | -2.13 (6.27) |
| Backslope | 4.77 (5.31) | 2.90 (5.43) | 3.93 (6.27) |
| Footslope | 10.24 (5.31) | 3.00 (5.43) | 3.54 (6.27) |
| Toeslope | -0.29 (5.31) | -8.45 (5.43) | -9.85 (6.27) |
| Stream | **18.43 * (9.01)** | 2.97 (9.21) | 4.90 (10.63) |
| $R^2$ | 0.07 | 0.07 | 0.08 |

**Table S3: Linear models predicting the relative abundance of classes within the universal compound pool.** Carbohydrates were not found in universal compounds and are omitted from this table. Values in cells are mean estimated effects ± standard error relative to the intercept, with the intercept expressed relative to zero. Bolded values were statistically significant at ***p < 0.001, **p < 0.01, *p < 0.05. For all, degrees of freedom = 67.

| | Lignins | Tannins | Condensed Hydrocarbons | Proteins | Amino Sugars | Lipids | Unsaturated Hydrocarbons |
|---|---|---|---|---|---|---|---|
| *Coefficient* | *Estimates* | | | | | | |
| Intercept: Site C1, 5 cm depth, shoulder position | **47.12 ***<br>(2.92)** | **5.62 ***<br>(0.47)** | **4.35 ***<br>(0.65)** | **1.37 ***<br>(0.21)** | **0.09 ***<br>(0.01)** | 0.12<br>(0.11) | **0.20 ***<br>(0.03)** |
| Site C2 | 2.93<br>(1.84) | -0.16<br>(0.30) | -0.22<br>(0.41) | 0.17<br>(0.13) | 0.00<br>(0.01) | 0.03<br>(0.07) | 0.04<br>(0.02) |
| Site H1 | 1.43<br>(1.84) | -0.03<br>(0.30) | **-0.96 *<br>(0.41)** | 0.08<br>(0.13) | -0.01<br>(0.01) | 0.03<br>(0.07) | 0.00<br>(0.02) |
| Site H2 | 0.96<br>(1.84) | 0.07<br>(0.30) | **-1.12 **<br>(0.41)** | -0.02<br>(0.13) | -0.01<br>(0.01) | 0.09<br>(0.07) | -0.01<br>(0.02) |
| 15 cm | 2.66<br>(1.90) | **-0.72 *<br>(0.31)** | **-1.11 *<br>(0.42)** | 0.08<br>(0.13) | -0.01<br>(0.01) | 0.02<br>(0.07) | 0.02<br>(0.02) |
| 30 cm | **10.41 ***<br>(1.90)** | **-1.28 ***<br>(0.31)** | **-2.85 ***<br>(0.42)** | **0.50 ***<br>(0.13)** | 0.01<br>(0.01) | **0.19 *<br>(0.07)** | **0.06 **<br>(0.02)** |
| 60 cm | **12.83 ***<br>(1.90)** | -0.59<br>(0.31) | **-3.36 ***<br>(0.42)** | **0.29 *<br>(0.13)** | -0.01<br>(0.01) | 0.04<br>(0.07) | 0.02<br>(0.02) |
| Backslope | 1.29<br>(1.90) | -0.04<br>(0.31) | -0.88 *<br>(0.42) | 0.21<br>(0.13) | 0.00<br>(0.01) | 0.12<br>(0.07) | -0.03<br>(0.02) |
| Footslope | 3.68<br>(1.90) | **-0.66 *<br>(0.31)** | **-1.90 ***<br>(0.42)** | **0.31 *<br>(0.13)** | -0.00<br>(0.01) | 0.03<br>(0.07) | **-0.05 *<br>(0.02)** |
| Toeslope | **7.14 ***<br>(1.90)** | -0.57<br>(0.31) | **-2.02 ***<br>(0.42)** | **0.42 **<br>(0.13)** | **0.02 *<br>(0.01)** | 0.08<br>(0.07) | -0.02<br>(0.02) |
| Stream | **11.37 ***<br>(3.22)** | -0.86<br>(0.52) | **-3.60 ***<br>(0.72)** | 0.31<br>(0.23) | -0.02<br>(0.01) | 0.05<br>(0.12) | -0.04<br>(0.04) |
| $R^2$ | 0.52 | 0.19 | 0.63 | 0.23 | 0.18 | 0.06 | 0.15 |

**Table S4: Linear models predicting the relative abundance of classes within the non-universal compound pool.** Values in cells are mean estimated effects ± standard error relative to the intercept, with the intercept expressed relative to zero. Bolded values were statistically significant at ***p < 0.001, **p < 0.01, *p < 0.05. For all, degrees of freedom = 67.

| *Coefficient* | Lignins | Tannins | Condensed Hydrocarbons | Proteins | Amino Sugars | Carbohydrates | Lipids | Unsaturated Hydrocarbons |
|---|---|---|---|---|---|---|---|---|
| Intercept: Site C1, 5 cm depth, shoulder position | **19.35 *** (1.38)** | **4.71 *** (0.58)** | **12.26 *** (2.52)** | **1.07 ** (0.37)** | **0.33 *** (0.08)** | **1.69 *** (0.30)** | **0.74 ** (0.23)** | **0.56 *** (0.07)** |
| Site C2 | -0.86 (0.87) | -0.55 (0.36) | -1.80 (1.59) | 0.19 (0.23) | -0.03 (0.05) | 0.15 (0.19) | 0.06 (0.14) | 0.04 (0.05) |
| Site H1 | **1.91 * (0.87)** | -0.35 (0.36) | -2.72 (1.59) | 0.42 (0.23) | 0.05 (0.05) | 0.01 (0.19) | 0.20 (0.14) | -0.01 (0.05) |
| Site H2 | **3.00 ** (0.87)** | -0.26 (0.36) | **-3.55 * (1.59)** | 0.44 (0.23) | **0.10 * (0.05)** | 0.37 (0.19) | 0.05 (0.14) | -0.04 (0.05) |
| 15 cm | 1.58 (0.90) | -0.45 (0.38) | -2.13 (1.64) | -0.26 (0.24) | **-0.18 *** (0.05)** | 0.24 (0.20) | 0.11 (0.15) | **0.14 ** (0.05)** |
| 30 cm | **2.52 ** (0.90)** | **-2.08 *** (0.38)** | **-9.39 *** (1.64)** | 0.39 (0.24) | **-0.15 ** (0.05)** | **0.80 *** (0.20)** | **0.60 *** (0.15)** | **0.27 *** (0.05)** |
| 60 cm | **3.27 *** (0.90)** | **-2.03 *** (0.38)** | **-10.89 *** (1.64)** | -0.28 (0.24) | **-0.20 *** (0.05)** | **0.67 ** (0.20)** | 0.16 (0.15) | **0.15 ** (0.05)** |
| Backslope | 0.95 (0.90) | -0.26 (0.38) | -1.84 (1.64) | 0.44 (0.24) | 0.07 (0.05) | -0.13 (0.20) | 0.29 (0.15) | -0.09 (0.05) |
| Footslope | **3.67 *** (0.90)** | **-1.01 ** (0.38)** | **-5.14 ** (1.64)** | **0.68 ** (0.24)** | 0.05 (0.05) | 0.27 (0.20) | **0.34 * (0.15)** | **-0.10 * (0.05)** |
| Toeslope | **2.47 ** (0.90)** | **-1.75 *** (0.38)** | **-7.33 *** (1.64)** | **0.97 *** (0.24)** | 0.10 (0.05) | 0.19 (0.20) | **0.61 *** (0.15)** | -0.09 (0.05) |
| Stream | **5.49 *** (1.52)** | **-1.92 ** (0.64)** | **-11.36 *** (2.78)** | 0.01 (0.41) | **-0.22 * (0.08)** | 0.63 (0.33) | 0.35 (0.25) | 0.02 (0.08) |
| $R^2$ | 0.43 | 0.50 | 0.56 | 0.27 | 0.29 | 0.24 | 0.30 | 0.32 |

**Table S5: Linear models predicting the intensity-weighted molecular mass of classes within the non-universal compound pool.** Values in cells are mean estimated effects ± standard error relative to the intercept, with the intercept expressed relative to zero. Bolded values were statistically significant at ***p < 0.001, **p < 0.01, *p < 0.05. For all, degrees of freedom = 67.

| *Coefficient* | Lignins | Tannins | Condensed Hydrocarbons | Proteins | Amino Sugars | Carbohydrates | Lipids | Unsaturated Hydrocarbons |
|---|---|---|---|---|---|---|---|---|
| Intercept: Site C1, 5 cm depth, shoulder position | **519.14 *** (12.17)** | **471.46 *** (15.51)** | **447.88 *** (18.92)** | **473.10 *** (16.79)** | **417.61 *** (19.66)** | **430.64 *** (18.47)** | **342.81 *** (12.66)** | **282.17 *** (10.87)** |
| Site C2 | -6.26 (7.70) | -5.01 (9.81) | -8.30 (11.97) | -12.18 (10.62) | -7.81 (12.44) | -0.34 (11.68) | -4.23 (8.01) | -1.89 (6.88) |
| Site H1 | **-18.34 * (7.70)** | 7.16 (9.81) | -10.67 (11.97) | **-30.42 ** (10.62)** | -9.44 (12.44) | 16.19 (11.68) | **-27.28 ** (8.01)** | 4.53 (6.88) |
| Site H2 | **-18.49 * (7.70)** | 3.83 (9.81) | -2.27 (11.97) | -20.47 (10.62) | 3.60 (12.44) | **28.44 * (11.68)** | **-18.83 * (8.01)** | 10.69 (6.88) |
| 15 cm | -2.29 (7.94) | 7.15 (10.11) | -9.57 (12.34) | 19.18 (10.94) | 9.34 (12.82) | 21.62 (12.04) | **27.93 ** (8.25)** | 9.84 (7.09) |
| 30 cm | **-17.56 * (7.94)** | -10.52 (10.11) | **-36.71 ** (12.34)** | -21.29 (10.94) | **-28.73 * (12.82)** | **54.31 *** (12.04)** | **34.96 *** (8.25)** | 13.64 (7.09) |
| 60 cm | -8.40 (7.94) | 7.94 (10.11) | 1.11 (12.34) | 6.78 (10.94) | 0.01 (12.82) | **72.43 *** (12.04)** | **31.79 *** (8.25)** | **36.91 *** (7.09)** |
| Backslope | -1.24 (7.94) | 5.09 (10.11) | 11.30 (12.34) | -4.02 (10.94) | 3.46 (12.82) | 20.29 (12.04) | 15.70 (8.25) | **12.45 (7.09)** |
| Footslope | -8.12 (7.94) | 12.82 (10.11) | 3.65 (12.34) | **-24.83 * (10.94)** | 1.77 (12.82) | **53.12 *** (12.04)** | 4.87 (8.25) | **26.47 *** (7.09)** |
| Toeslope | **-25.69 ** (7.94)** | -0.23 (10.11) | -8.91 (12.34) | **-61.28 *** (10.94)** | -18.63 (12.82) | **65.37 *** (12.04)** | 4.96 (8.25) | **21.86 ** (7.09)** |
| Stream | -11.00 (13.46) | 18.84 (17.14) | 12.18 (20.92) | 9.43 (18.55) | 16.31 (21.73) | **102.10 *** (20.42)** | **47.08 ** (13.99)** | **48.16 *** (12.02)** |
| $R^2$ | 0.21 | -0.02 | 0.10 | 0.46 | 0.10 | 0.55 | 0.33 | 0.39 |

**Table S6: Linear models predicting changes in environmental variables at each depth and hillslope position at each site.** Values in cells are mean estimated effects on a log-scale ± standard error relative to the intercept, with the intercept expressed relative to zero. Bolded values were statistically significant at ***p < 0.001, **p < 0.01, *p < 0.05. For all, degrees of freedom = 67.

| *Coefficient* | DOC (mg $L^{-1}$) | Bacterial Productivity ($10^{-3}$ mg C $L^{-1}$) | Humic Substances: Low Molecular Weight Substances | Carbon: Nitrogen (Total) |
|---|---|---|---|---|
| Intercept: Site C1, 5 cm depth, shoulder position | **4.98 *** (1.23)** | **0.01 *** (0.01)** | **2.33 * (0.80)** | **22.20 *** (7.16)** |
| Site C2 | 1.00 (0.13) | NA | 1.12 (0.21) | 0.72 (0.12) |
| Site H1 | **0.51 *** (0.07)** | **2.40 * (0.95)** | **0.60 ** (0.11)** | **0.33 *** (0.06)** |
| Site H2 | **0.52 *** (0.07)** | NA | 0.74 (0.14) | **0.23 *** (0.04)** |
| 15 cm | 0.79 (0.11) | 0.55 (0.30) | **1.50 * (0.28)** | **0.66 * (0.11)** |
| 30 cm | **0.48 *** (0.06)** | 0.38 (0.20) | 0.91 (0.17) | **0.56 ** (0.10)** |
| 60 cm | **0.28 *** (0.04)** | **0.12 ** (0.07)** | 0.97 (0.19) | **0.46 *** (0.08)** |
| Backslope | **0.74 * (0.10)** | 2.70 (1.41) | 0.86 (0.16) | 0.83 (0.14) |
| Footslope | **0.33 *** (0.04)** | **4.83 ** (2.63)** | **0.51 *** (0.09)** | **0.66 * (0.12)** |
| Toeslope | **0.20 *** (0.03)** | **0.02 *** (0.01)** | **0.39 *** (0.08)** | 0.80 (0.14) |
| Stream | **8.40 *** (2.15)** | NA | **2.20 * (0.78)** | 1.75 (0.58) |
| $R^2$ | 0.84 | 0.78 | 0.42 | 0.63 |

**Table S7: Differentially expressed CAZymes along the depth gradient.** For each CAZyme, we calculated the mean number of normalised reads estimated over all the samples, along with the mean ± standard error (SE) of the effect of soil depth estimated from negative binomial generalized linear models using the R package DESeq2[8]. P-values from a Wald test were corrected for multiple comparisons with a Benjamini-Hochberg adjustment.

| Carbohydrate-active enzyme | Mean count | Mean effect size | SE effect size | Adjusted p-value |
|---|---|---|---|---|
| Carbohydrate-Binding Module Family 8 | 96.21 | -2.60 | 0.54 | 3.35 E-04 |
| Polysaccharide Lyase Family 11 | 284.54 | -0.53 | 0.11 | 3.78 E-04 |
| Polysaccharide Lyase Family 1, Subfamily 2 | 910.32 | -1.63 | 0.39 | 2.94 E-03 |
| Glycoside Hydrolase Family 13 Subfamily 21 | 217.85 | -0.95 | 0.24 | 5.53 E-03 |
| Glycoside Hydrolase Family 5, Subfamily 54 | 91.11 | -1.99 | 0.51 | 5.53 E-03 |
| Glycoside Hydrolase Family 116 | 1476.49 | -0.99 | 0.26 | 8.03 E-03 |
| Glycoside Hydrolase Family 64 | 155.07 | 1.41 | 0.41 | 2.21 E-03 |
| Glycoside Hydrolase Family 65 | 3457.61 | -0.76 | 0.22 | 2.38 E-02 |
| Carbohydrate Esterase Family 3 | 609.33 | 0.58 | 0.18 | 3.31 E-02 |
| Glycoside Hydrolase Family 6 | 232.70 | 1.20 | 0.37 | 3.31 E-02 |
| Carbohydrate-Binding Module Family 12 | 242.67 | -1.08 | 0.34 | 3.78 E-02 |
| Glycoside Hydrolase Family 20 | 4213.18 | -0.57 | 0.18 | 3.78 E-02 |
| Glycoside Hydrolase Family 17 | 950.16 | 0.68 | 0.22 | 3.40 E-02 |
| Glycoside Hydrolase Family 73 | 192.13 | 0.54 | 0.17 | 3.40 E-02 |
| Carbohydrate-Binding Module Family 48 | 1538.12 | -1.28 | 0.42 | 4.18 E-02 |
| Carbohydrate-Binding Module Family 6 | 750.91 | 1.53 | 0.51 | 4.58 E-02 |
| Auxiliary Activity Family 2 | 3419.56 | 1.21 | 0.41 | 4.78E-02 |

**Table S8: Differentially expressed CAZymes along the hillslope gradient.** For each CAZyme, we calculated the mean number of normalised reads estimated over all the samples, along with the mean ± standard error (SE) of the effect of soil depth estimated from negative binomial generalized linear models using the R package DESeq2[8]. P-values from a Wald test were corrected for multiple comparisons with a Benjamini-Hochberg adjustment. Genes explaining at least 2.5% of the variation in compound class are bolded.

| Carbohydrate-active enzyme | Mean count | Mean effect size | SE effect size | Adjusted p-value |
|---|---|---|---|---|
| **Auxiliary Activity 001** | **32916.22** | **0.93** | **0.31** | **4.24E-24** |
| **Polysaccharide Lyase 001, Subfamily 2** | **910.32** | **1.28** | **0.39** | **1.18E-11** |
| **Glycoside Hydrolase Family 13, Subfamily 26** | **4268.76** | **-0.49** | **0.16** | **2.97E-10** |
| **Glycoside Hydrolase Family 13, Subfamily 18** | **335.14** | **1.02** | **0.34** | **3.90E-07** |
| **Glycoside Hydrolase Family 51** | **5063.26** | **1.07** | **0.30** | **1.44E-06** |
| **Glycoside Hydrolase Family 135** | **348.96** | **1.34** | **0.42** | **1.53E-05** |
| Glycoside Hydrolase Family 153 | 220.19 | 2.31 | 0.22 | 5.84E-05 |
| Glycoside Hydrolase Family 127 | 2806.18 | 1.51 | 0.20 | 1.18E-04 |
| Glycoside Hydrolase Family 44 | 5530.20 | 1.96 | 0.28 | 2.33E-04 |
| Glycoside Hydrolase Family 125 | 398.41 | 1.90 | 0.32 | 2.57E-04 |
| Glycosyltransferase Family 30 | 5307.16 | 1.13 | 0.20 | 2.57E-04 |
| Glycosyltransferase Family 111 | 14.38 | 4.55 | 0.88 | 2.69E-04 |
| Glycosyltransferase Family 21 | 49598.10 | 3.05 | 0.62 | 3.23E-04 |
| Glycoside Hydrolase Family 65 | 3457.61 | -1.05 | 0.22 | 3.58E-04 |
| Polysaccharide Lyase Family 9, Subfamily 1 | 97.78 | -1.31 | 0.29 | 3.58E-04 |
| Glycoside Hydrolase Family 13, Subfamily 16 | 4526.45 | -0.44 | 0.10 | 3.58E-04 |
| Glycoside Hydrolase Family 5, Subfamily 7 | 329.90 | 0.87 | 0.19 | 1.02E-03 |
| Glycoside Hydrolase Family 29 | 9898.17 | 1.66 | 0.37 | 1.34E-03 |
| Glycoside Hydrolase Family 146 | 144183.44 | 3.32 | 0.75 | 1.42E-03 |
| Carbohydrate-Binding Module Family 9 | 6125.91 | 1.43 | 0.33 | 1.42E-03 |
| Glycosyltransferase Family 17 | 71.84 | 1.89 | 0.44 | 1.44E-03 |
| Glycosyltransferase Family 5 | 10438.03 | 0.91 | 0.21 | 2.13E-03 |
| Glycosyltransferase Family 22 | 353.05 | -0.72 | 0.17 | 4.77E-03 |
| Glycoside Hydrolase Family 3 | 28555.42 | 0.82 | 0.20 | 5.36E-03 |
| Glycoside Hydrolase Family 140 | 5303.31 | 2.23 | 0.56 | 5.36E-03 |
| Polysaccharide Lyase Family 6 | 62.10 | -2.27 | 0.57 | 6.83E-03 |
| Carbohydrate-Binding Module Family 50 | 3115.27 | -0.35 | 0.09 | 7.02E-03 |
| Glycoside Hydrolase Family 12 | 120.38 | -0.67 | 0.17 | 7.02E-03 |
| Glycoside Hydrolase Family 5, Subfamily 54 | 91.11 | -1.88 | 0.52 | 7.18E-03 |
| Glycoside Hydrolase Family 30, Subfamily 7 | 35.34 | -1.18 | 0.33 | 9.47E-03 |
| Polysaccharide Lyase Family 14, Subfamily 3 | 107.70 | 0.90 | 0.26 | 9.59E-03 |

| | | | | |
|---|---|---|---|---|
| Glycosyltransferase Family 107 | 91.25 | 0.81 | 0.23 | 1.03E-02 |
| Polysaccharide Lyase Family 25 | 64.03 | -3.05 | 0.87 | 1.11E-02 |
| Glycosyltransferase Family 84 | 4586.90 | 0.86 | 0.25 | 1.13E-02 |
| Glycoside Hydrolase Family 78 | 4423.06 | 0.61 | 0.18 | 1.17E-02 |
| Glycoside Hydrolase Family 5, Subfamily 10 | 76.25 | -0.63 | 0.19 | 1.17E-02 |
| Glycoside Hydrolase Family 63 | 5970.43 | 0.85 | 0.25 | 1.19E-02 |
| Glycoside Hydrolase Family 106 | 1733.42 | 0.69 | 0.21 | 1.44E-02 |
| Glycoside Hydrolase Family 141 | 857.12 | 1.10 | 0.33 | 1.70E-02 |
| Auxiliary Activity Family 3 | 42840.58 | 1.27 | 0.38 | 1.77E-02 |
| Carbohydrate-Binding Module Family 48 | 1538.12 | -1.37 | 0.42 | 1.95E-02 |
| Glycoside Hydrolase Family 67 | 654.10 | -0.32 | 0.10 | 1.95E-02 |
| Glycoside Hydrolase Family 92 | 708.76 | 0.42 | 0.13 | 1.99E-02 |
| Glycoside Hydrolase Family 37 | 2171.48 | 1.28 | 0.41 | 2.05E-02 |
| Polysaccharide Lyase Family 11 | 284.54 | -0.36 | 0.12 | 2.18E-02 |
| Glycoside Hydrolase Family 144 | 3152.45 | 0.46 | 0.15 | 2.18E-02 |
| Glycoside Hydrolase Family 16 | 130.03 | -0.54 | 0.18 | 2.19E-02 |
| Glycosyltransferase Family 4 | 120988.48 | 0.42 | 0.14 | 2.39E-02 |
| Glycosyltransferase Family 7 | 40.61 | 1.25 | 0.42 | 2.66E-02 |
| Glycosyltransferase Family 39 | 3342.74 | -0.57 | 0.19 | 2.74E-02 |
| Glycoside Hydrolase Family 87 | 306.14 | 0.91 | 0.31 | 3.05E-02 |
| Carbohydrate-Binding Module Family 47 | 241.00 | 1.31 | 0.46 | 3.30E-02 |
| Glycoside Hydrolase Family 43, Subfamily 12 | 137.27 | -0.60 | 0.21 | 3.61E-02 |
| Glycoside Hydrolase Family 13, Subfamily 13 | 401.51 | -0.45 | 0.16 | 3.97E-02 |
| Glycoside Hydrolase Family 47 | 1117.19 | -1.15 | 0.41 | 3.97E-02 |
| Glycoside Hydrolase Family 48 | 181.55 | 0.56 | 0.20 | 3.97E-02 |
| Glycoside Hydrolase Family 5, Subfamily 17 | 54.05 | -0.48 | 0.17 | 3.99E-02 |
| Glycoside Hydrolase Family 43, Subfamily 28 | 103.16 | -0.71 | 0.26 | 4.03E-02 |
| Glycoside Hydrolase Family 5, Subfamily 1 | 154.20 | -0.48 | 0.17 | 4.06E-02 |
| Glycoside Hydrolase Family 133 | 5371.77 | -0.48 | 0.18 | 4.13E-02 |
| Carbohydrate-Binding Module Family 57 | 1376.46 | 0.52 | 0.19 | 4.35E-02 |

**Table S9: Data sources used to generate Figure 5.** The mean percentage of molecules shared with a deep-sea reference sample was calculated for different sample types. If there was more than one study for a given sample type, the number of samples from each study is given in parenthesis.

| Sample Type | Number of Samples | Study |
|---|---|---|
| 5 cm Depth (Soil) | 19 | this one, Simon et al.[9] (3) |
| 15 cm Depth (Soil) | 20 | this one, Simon et al.[9] (4) |
| 30 cm Depth (Soil) | 17 | this one, Simon et al.[9] (1) |
| 60 cm Depth (Soil) | 17 | this one, Simon et al.[9] (1) |
| Shoulder (Soil) | 16 | this one |
| Backslope (Soil) | 16 | this one |
| Footslope (Soil) | 16 | this one |
| Toeslope (Soil) | 16 | this one |
| Stream | 15 | this one (4), Hutchins et al. 2017[10] (11) |
| River | 144 | Hutchins et al. 2017[10] |
| Lake | 116 | Kellerman et al. 2014[11](115) Zark & Dittmar 2018[12] (1) |
| Bog | 4 | Simon et al. 2018[9] |
| Sea Surface | 4 | Zark & Dittmar 2018[12] |
| Aquifer | 2 | Simon et al. 2018[9] |
| Deep-sea | 4 | Simon et al. 2018[9] |

**Table S10: Summary of physical and chemical environmental variables measured in soil pore water.** D.L. = Detection limit.

| Parameter/Range/ Detection Limit | Procedure | Principal Equipment |
|---|---|---|
| pH (3 - 9) | Orion Ross Ultra glass Electrode | Man-Tech PC-Titrate, Orion Thermo combination pH electrode |
| Specific Conductance (0 - 150 umho) | PCE-96-CT1003 electrode us/cm @ 25 C | Man-Tech PC-Titrate, with 4510 Conductivity meter |
| Total Alkalinity (0 - 2 meq/l ) | Electrometric Titration | Man-Tech PC-Titrate, Titra-Sip titrator, Orion Thermo combination pH electrode |
| Total Nitrogen (0 - 2 ppm) D.L. - 0.05ppm | Automated Cadmium Reduction | Technicon Autoanalyser II - NO2+ NO3 channel - Autoclave Digestion - N.A.P. software |
| NH4 as N (0 - 500ppb ) D.L. - 10ppb | Automated Sodium Nitroprusside, Filtered .45um | Seal Analytical AA3 Autoanalyzer AACE 6.07 software |
| NO2+NO3 as N (0 - 2ppm) D.L. - .04ppm | Automated Cadmium Reduction, Filtered .45um | Seal Analytical AA3 Autoanalyzer AACE 6.07 software |
| Total Phosphorus (unfiltered) (0 - 60ppb) D.L. - 1ppb | Automated Molybdophosphoric Blue | Technicon Autoanalyser II - N.A.P. software - Autoclave Digestion |
| Potassium – low end accuracy (water) 0.01ppm | ICP-MS | Agilent 7700x Inductively Coupled Plasma Instrument - Masshunter software |
| Sodium - low end accuracy (water) 0.01ppm | ICP-MS | Agilent 7700x Inductively Coupled Plasma Instrument - Masshunter software |
| Calcium – low end accuracy (water) 0.01ppm | ICP-MS | Agilent 7700x Inductively Coupled Plasma Instrument - Masshunter software |
| Magnesium – low end accuracy (water) 0.01ppm | ICP-MS | Agilent 7700x Inductively Coupled Plasma Instrument - Masshunter software |
| Sulphate (0 - 10ppm) D.L. -.2ppm | Conductance - ion exchange suppression | Dionex ICS 1100 Ion Chromatograph-Chromeleon 7.0 software |
| Total Sulphur as Sulphate | ICP-MS | Agilent 7700x Inductively Coupled Plasma Instrument- Masshunter software |
| Chloride (0 - 5ppm) D.L. - .2 ppm | Conductance - Ion Exchange Suppression | Dionex ICS 1100 Ion Chromatograph-Chromeleon 7.0 software |
| Silica Oxide (0 - 50ppm) & (0-7ppm) D.L. - .25 ppm | Automated Ascorbic Acid | Technicon Autoanalyser II - N.A.P. software |

| | | |
|---|---|---|
| Iron - low end accuracy (water) 0.005ppm | ICP-MS | Agilent 7700x Inductively Coupled Plasma Instrument - Masshunter software |
| Aluminum – low end accuracy (water) 0.005ppm | ICP-MS | Agilent 7700x Inductively Coupled Plasma Instrument - Masshunter software |
| Manganese - low end accuracy (water) 0.0005ppm | ICP-MS | Agilent 7700x Inductively Coupled Plasma Instrument - Masshunter software |
| Zinc – low-end accuracy (water) 0.001ppm | ICP-MS | Agilent 7700x Inductively Coupled Plasma Instrument - Masshunter software |
| Copper - low-end accuracy (water) 0.0005ppm | ICP-MS | Agilent 7700x Inductively Coupled Plasma Instrument - Masshunter software |
| Nickel – low-end accuracy (water) 0.0005ppm | ICP-MS | Agilent 7700x Inductively Coupled Plasma Instrument - Masshunter software |
| Cadmium – low-end accuracy (water) 0.0005ppm | ICP-MS | Agilent 7700x Inductively Coupled Plasma Instrument - Masshunter software |
| Lead – low-end accuracy (water) 0.0005ppm | ICP-MS | Agilent 7700x Inductively Coupled Plasma Instrument - Masshunter software |
| Dissolved Organic Carbon (0 - 20 ppm)<br>DL. - .4ppm | Sample Purged - N2 Acid & Potassium Persulphate - U.V. Rad. - Dialysis - O.C. Inversely Meas., Filtered .45um | Seal Analytical AA3 Autoanalyzer - AACE 6.07 software |
| Dissolved Inorganic Carbon (0 - 5ppm) DL. - .5ppm | Sample Acidified H2SO4 - CO2 Dialysis Thru Membrane - I.C. Inversely Meas., Filtered .45um | Seal Analytical AA3 Autoanalyzer - AACE 6.07 software |
| Soluble Reactive Phosphorus (0 - 60ppb)<br>D.L. - 1ppb | Automated Molybdophosphoric Blue, Filtered .45um | Technicon Autoanalyser II - N.A.P. software |
| High-level NH4 as N<br>(0 – 100ppm )<br>D.L. – 2ppm | Automated Sodium Nitroprusside | Technicon Autoanalyser II - N.A.P. software |
| Arsenic – low-end accuracy (water) 0.0005ppm | ICP-MS | Agilent 7700x Inductively Coupled Plasma Instrument - Masshunter software |
| Cobalt – low-end accuracy (water) 0.0005ppm | ICP-MS | Agilent 7700x Inductively Coupled Plasma Instrument - Masshunter software |
| Chromium – low end accuracy (water) 0.0005ppm | ICP-MS | Agilent 7700x Inductively Coupled Plasma Instrument - Masshunter software |
| Selenium – low end accuracy (water) 0.001ppm | ICP-MS | Agilent 7700x Inductively Coupled Plasma Instrument - Masshunter software |
| Strontium – low-end accuracy (water) 0.001ppm | ICP-MS | Agilent 7700x Inductively Coupled Plasma Instrument - Masshunter software |

### Supplementary Methods

***Rarefaction of DOM:*** We combined published data from Table S9 into a sample × molecular formula matrix. We then randomly generated 10,000 intervals between 250 to 20078, and randomly selected the corresponding molecular formula from the sample set. We then plotted the proportion of universal compounds in each sample as a function of the total number of sampled formulae, repeating this process 999 times. We visually identified a threshold of 6000 molecular formulae, after which additional sampling did not change the proportion of universal compounds per sample.

***Bacterial productivity:*** In the laboratory, 1 g of soil from each sample was mixed with 50 mL distilled water using a gyratory shaker (200 rev $min^{-1}$) and soil particles were removed by centrifugation (1000 *g*) for 10 min. We then added 3.74 µL of $^3$H-labeled leucine (1 mCi $mL^{-1}$, PerkinElmer, USA) to 1.5 mL of bacterial suspension. Blanks were prepared by immediately adding 75 µL of 100% trichloroacetic acid (TCA), killing any live cells, and providing a measure of background $^3$H leucine incorporation that was later subtracted from the incubated values. After 2 h of incubation at room temperature (~22°C), each sample received 75 µL of ice-cold 100% trichloroacetic acid (TCA) to stop incorporation. Samples were subsequently centrifuged at 13,000 *g* for 10 min and the supernatant was discarded. Each sample was then washed once with 5% TCA, 1.5 mL ice-cold 80% ethanol, and 0.2 mL 1M NaOH. Samples were vortexed and maintained at 90°C for 1 h. After cooling to room temperature, we added 1 mL of Ultima Gold scintillation fluid (Perkin Elmer, USA) to each sample and vortexed for 30 s. $^3$H incorporation was measured with a LS6500 liquid scintillation counter (Beckman Coulter, USA).

***Bacterial cell counts:*** Each sample was defrosted and we added 1.2 mL of a detergent: 250 mM tetrasodium pyrophosphate [TSP; pH 8.0] containing Tween 80, 0.5% final concentration. The solution was then vortexed for 30 s and shaken for 2 hr at 4°C. From the resulting slurry, 1 mL was layered slowly onto 0.5 mL of Histodenz solution (80% [wt/vol] prepared in 50 mM sterile TSP buffer). Bacterial cells and soil particles were separated by high-speed centrifugation (14,000 *g*) for 30 min. The upper and middle cell-containing phases (including the thin layer on top of the Histodenz cushion liquid phase) were carefully recovered and mixed with 1 mL of 50 mM TSP buffer and centrifuged at 17,000 *g* for 25 min. The supernatant was removed, and the cell pellet containing the cell fraction was resuspended in 0.8 ml of 50mM TSP buffer. The pellet slurries were vortexed and then stained with SYBR Green (final concentration 1×) and incubated for 20 min at room temperature in the dark. Samples were vortexed again before cytometry counts, and between each sample, 100 µL of sterile TSP buffer was passed through the flow cytometer.

***Principal coordinates of neighbour matrices:*** The distance between the lysimeters was represented as a Euclidean distance matrix. Distances were calculated from the average latitude and longitude of the three replicate lysimeters at the same hillslope position. The resulting PCNM decomposes any spatial relationships between sample locations at the same site. This captures distinct information which justifies its inclusion alongside the ordinal variables (depth) and categorical variables (hillslope). We did not compute distances between lysimeters in different catchments by setting a threshold distance of 500 m[13].
